# Parallel Sampling of Protein-Ligand Dynamics

**DOI:** 10.1101/2024.07.08.602465

**Authors:** Matthew R. Masters, Amr H. Mahmoud, Markus A. Lill

## Abstract

Molecular dynamics (MD) simulations of protein-ligand complexes are essential for computer-aided drug design. In particular they enable the calculation of free energies and thus binding affinities. However, these simulations require significant computational resources and can take days to weeks to achieve relatively short timescales compared to biologically relevant timescales. To address this issue, we introduce a method for non-sequential generation of MD samples using a generative deep neural network trained on a large corpus of protein-ligand complex simulations. The method generates accurate protein-ligand complexes with full protein and ligand flexibility and is able to recapitulate the conformation space sampled by MD simulations with high coverage. This development is a step forward towards one-shot molecular sampling that can be utilized in the calculation of protein-ligand free energies.^3^

## 1 Introduction

Molecular dynamics (MD) simulations have long been a pivotal tool in the field of computational chemistry and biophysics, allowing for the probing of molecular interactions and dynamics at an atomic level. They provide invaluable insights into the behaviors and characteristics of systems that are often unattainable through experimental methodologies alone. Notably, they play an instrumental role in predicting the behaviors of a broad range of chemical and biochemical systems, from protein folding dynamics to material properties at the nanoscale.

Simulations of protein-ligand complexes are particularly critical in the fields of drug design. For example, they enable the estimation of free energies associated with ligand binding, a key factor in optimizing lead candidates [1, 2, 3]. Using this information, researchers can predict how modifications to ligand structures influence binding characteristics. This ability to predict and optimize binding affinity before synthesis is invaluable, dramatically reducing the cost and increasing the efficiency of developing new therapeutics [4, 5]. Protein-ligand simulations can also be used to better understand the binding and unbinding dynamics of drugs [6], induced conformational changes [7, 8], and to determine the stability of a particular pose [9, 10]. These tasks are especially useful for modelled complexes that come from molecular docking, since they often rely upon a single, static structure which cannot be rigorously evaluated unless simulated using dynamics.

However, the predominant issue with MD simulations is their computational and time requirements. Despite the transformative insights they offer, MD simulations demand significant computational resources due to the complexity and scale of the calculations involved. Exhaustive sampling of the conformational space is limited due to the rugged, high-dimensional energy landscape, which requires the evaluation of forces and movements for potentially hundreds of thousands of atoms over millions of time steps. This computational intensity necessitates the use of high-performance computing (HPC) systems equipped with powerful processors and often, parallel computing architectures [11]. Furthermore, the need for extensive simulation times to achieve statistically significant results can result in high energy consumption and substantial operational costs. Advances in computational methods have sought to address these challenges by reducing the time required to reach convergence [12, 13]. However, sampling must still be done sequentially and simulations can still take days to even months to complete.

To address this shortcoming, we introduce a generative AI method for non-sequential generation of MD samples using a deep neural network architecture. The model utilizes a diffusion-based concept trained on a corpus of over a million frames from a large, diverse set of molecular dynamics simulations. We show that the trained model is capable of sampling unseen protein-ligand systems with great coverage compared to the MD simulation samples. This represents a step forward towards one-shot sampling of protein-ligand systems which has the potential to enable ultra-fast, accurate free energy estimations adhering to statistical mechanics principles.

## 2 Background and Related Work

Molecular dynamics (MD) simulations are integral to computational biology, providing a dynamic picture of molecular interactions at an atomic level [14, 15]. By simulating the motions of atoms and molecules, MD allows researchers to capture the complex interplay between components in a biological system, which static experimental approaches like X-ray crystallography or cryo-electron microscopy cannot achieve [16, 17]. This dynamic insight is essential for elucidating the mechanisms of biomolecular functions and the structural changes associated with protein-ligand binding [18, 19, 20].

The ability of MD simulations to provide these insights has dramatically improved with advances in computational techniques and the increased availability of high-performance computing resources [21, 22, 23, 24]. These developments have extended the scale and duration of simulations, enhancing their accuracy and the depth of biological phenomena they can model. Additionally, the integration of MD data with predictive modeling techniques, such as deep learning, has further expanded their applicability in drug discovery, allowing for more accurate predictions of molecular behavior [25, 26, 27].

### 2.1 Diffusion Models for Molecular Sampling

Diffusion models have emerged as an effective deep learning model to sample molecular systems. Many molecular diffusion models have recently been introduced, for example in the applications of protein structure prediction [28], de novo ligand generation [29], learning molecular force fields [30], and molecular docking [31, 32, 33, 34, 35]. The application of diffusion models to protein-ligand systems began with the seminal work of DiffDock [36]. The approach has since been adapted and developed into a number of different papers. For example, DiffDock was adapted to work on user-defined pockets [37], and DynamicBind adapted it to have full protein flexibility, allowing for large induced conformational changes [31]. Integration of physics priors showed additional improvement in generalization power of diffusion models [38]. Our approach takes significant influence from the work of DiffDock and DynamicBind and utilizes a similar SE(3)-equivariant network architecture. More recently, the works of RoseTTAFold and AlphaFold have released new versions which migrate to a diffusion-based architecture that is capable of modelling arbitrary molecular structures with breakthrough accuracy [33, 39, 40, 41]. In addition, there are related flow matching methods with similar applications to molecules [42, 43, 44], which resemble the interpolation-based method we present. While all these results are impressive steps forward in the utilization of deep learning models for molecular modelling, they only train using static crystal structures and do not leverage the rich dynamics of MD.

## 3 Materials and Methods

### 3.1 Diffusion-based Model

Our model employs a diffusion-based architecture, which is pivotal for learning the dynamic transitions between molecular conformations as derived from molecular dynamics simulations. Diffusion models are typically divided into two phases: the forward and reverse diffusion processes. The forward diffusion process starts with real samples from the target data distribution and is progressively noised over many steps. This process simulates a Markovian trajectory where each step represents an increasingly distorted version of the original structure. The reverse diffusion process aims to restore the original molecular configuration by sequentially reducing the noise and thereby refining the structure. By training the model to effectively reverse the noise addition, it learns to trace back the most probable path to the native state of the protein-ligand complex.

Under the typical formulation of diffusion models, the data is noised until it becomes identical to the normal distribution, removing all structure from the original data. In contrast to this approach, our model learns the interpolation between two frames without the addition of noise (See Figure 1). This is advantageous because it allows us to initialize from the same starting frame as the MD simulation and the model only has to learn the transition from one physically-meaningful frame to the next. With conventional diffusion models, samples are initialized from the normal distribution and progressively denoised using the diffusion model. This so-called “black-hole initialization” means that the diffusion model must accurately model both structured and unstructured data at various scales. Using our approach, the model receives a well structured input already and only has to make small perturbations to simulate going from one frame to another.

**Figure 1:**
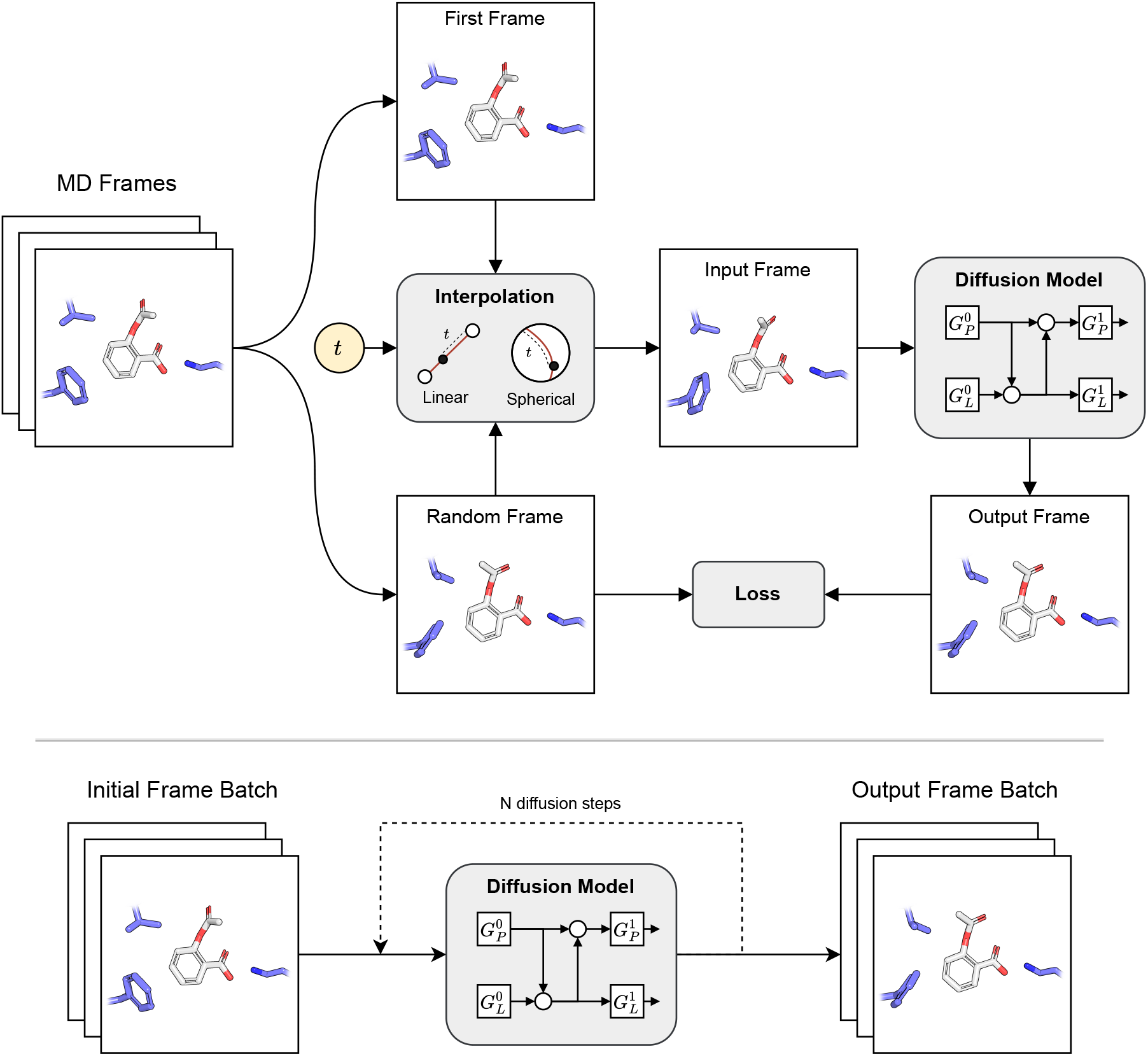
Overview of the training (top) and inference (bottom) procedures. During training, rigid interpolation is used to generate an intermediate state between the initial frame and a randomly sampled one. The interpolated frame is provided to the equivariant diffusion model which attempts to predict the sampled frame.

To this end we utilized interpolation between a starting frame and another frame from the simulation selected at random. The interpolation from a starting frame to the sampled frame is done through a combination of linear and spherical interpolation. Since the *C*_*α*_ and ligand centroid positions are simply Cartesian coordinates, we use linear interpolation to go between these variables of the two frames. Similarly, we use linear interpolation for torsion angles of protein and ligand, with special handling for the torus topology. However, rigid rotations of the ligand and residues are less straight-forward and require spherical linear interpolations (SLERP) in order to transform along the arc of sphere [45].

The interpolated frame can then be encoded into protein and ligand graphs as input for the network. Let *𝒢*_*L*_ = (*𝒱*_*L*_, *ℰ*_*L*_) be a graph of the ligand with vertices (*𝒱*_*L*_ = (**h**_*i*_, **x**_*i*_) representing heavy atoms and edges *ℰ*_*L*_ representing nearby or covalently bound atoms. The edges are featurized using the bond type (single, double, triple, or aromatic) and Euclidean distance in Ångstrom. Similarly, the protein is encoded into graph *𝒢*_*P*_ = (*𝒱*_*P*_, *ℰ*_*P*_) where nodes represent the *C*_*α*_ positions and edges represent Euclidean distances between nearby residues. The protein nodes are featurized using a combination of residue type, pre-trained large language sequence embeddings from ESM [46], and side-chain torsion angles expressed as zero-padded scalar features. These angles include five flexible chi angles and two symmetric chi angles, converted into sine and cosine values for uniqueness. Additionally, the orientation of the protein backbone is captured by two unit vectors representing normalized directional vectors between specific backbone atoms, defining a local rigid frame.

The diffusion model incorporates SE(3)-transformer layers that serve to interact and update the protein and ligand representations. Each layer updates both the protein and ligand graph representations by first updating the ligand based on the protein and then protein based on the ligand followed by self-attention. These layers are SE(3)-equivariant and respect rotation, translation, and reflection symmetries by making use of tensor products of irreducible representations [36, 47]. Messages are constructed as the tensor product between spherical harmonic representations of each edge vector with the current node representation:

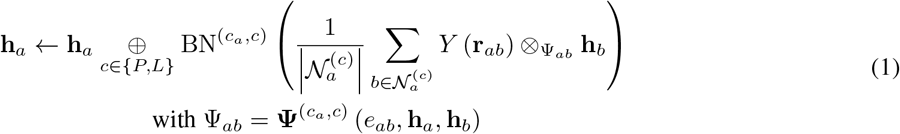

where **h**_*a*_ are the node features, **r**_*ab*_ is the Euclidean distance, and 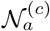 are the neighbors of node *a* with category *c* ∈ {*P, L*} (protein or ligand). The spherical harmonics are denoted as *Y* and BN represents the batch normalization. The learnable weights are contained in **Ψ**, which consists of a multi-layer perceptron (MLP) which takes the edge embeddings *e*_*ab*_ and features **h**_*a*_ and **h**_*b*_.

Following the last interaction layer, the node and edge embeddings are used to predict vectors which update the input conformation. For the ligand, this consists of vectors 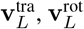 and 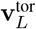 representing updates to the translation, rotation, and torsions of the ligand. The vectors are designated similarly for the protein. In order to calculate 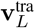 and 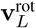, a convolution between ligand nodes and the ligand centroid is performed:

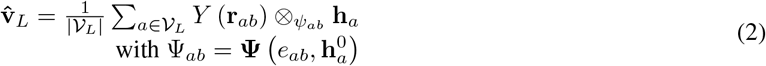

where *b* is the hypothetical centroid node. As with the interaction layers, **Ψ** represents an MLP that operate on the single node and edge features. Equation 2 is computed using separate weights for the translation and rotation vectors, yielding both 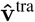 and 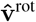. These vectors are then scaled using a single input MLP conditioned on a sinusoidal embedding *s*_*t*_ of the time *t* sampled earlier:

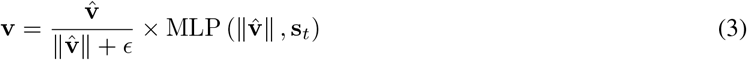

where *ϵ* is a small constant inserted for numerical stability at small 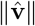. In order to predict the torsion updates for each rotatable bond, another convolution is performed between ligand nodes and a hypothetical node *c* placed at the center of the rotatable bond *b*:

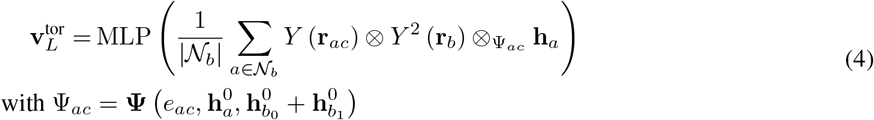

The calculation of the protein vectors 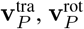 and 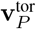 take a different form that does not rely on center points. They are calculated by simply passing the final interaction representation through a multi-layer perceptron 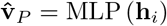 with different weights for vector. The translation and rotation vectors are then transformed by Equation 3 described earlier. The torsional updates have no further scaling 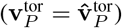.

Once the conformational updates have been predicted by the network, they can be applied to the protein and ligand. First, ligand torsion updates are applied and the resulting conformer is aligned to the previous conformer to prevent undue influence of torsions on translation or rotation movement. The ligand is rotated in place and then translated according to the predicted update. The protein is then updated by rotating and translating the frame, and propagating the torsional updates if necessary. The model can then be trained using a dataset of MD frames of protein-ligand complexes as shown in Figure 1. At each step, the model is trained to predict the translation, rotation, and torsional updates for the ligand and protein that would lead to the final “denoised” state. The predicted updates are compared with the true updates via the mean squared error loss with equal weighting.

### 3.2 Post-processing

Without any further processing, the model tends to produce poses that are slightly more explorative than those seen in MD frames. Additionally, occasionally the model will produce a pose that clashes with the protein surface or does not agree with the other samples. For this reason, we employed two post-processing steps that enhance the accuracy of the generated samples. First, we employ a steric clash filter which detects when a ligand pose clashes with the protein and removes this sample. To this end, we simply compute pairwise protein-ligand distances and detect when the distance is within the vdW radii plus some tolerance. Any clashing samples are removed and the inference process continues until there are the desired number of samples. Additionally, we applied outlier detection to the first two principle components in order to remove poses that were drastically different and did not agree with the rest of the samples. While these post-processing conditions are not ideal, we believe that further developments, particularly in adding physical biases to the model can alleviate the need for such filtering.

### 3.3 Dataset and Preparation

The PDBBind data provides 19,443 protein–ligand structures that have been experimentally determined and have affinity values available [48, 49]. Recently, a dataset containing molecular dynamics simulations of these complexes has been released [50]. The dataset, called MISATO, was rigorously curated to correct structural inaccuracies, including adjustments for protonation states and bond geometries. After some filtering, MD simulations of 16,972 protein–ligand complexes were performed. The dataset was randomly split, with a test set of 500 samples.

### 3.4 Training and Inference

Training was performed for 400 epochs with a learning rate of 0.001 with the Adam optimizer. Systems and frames were sampled uniformly from the training set and collated into batches of 32. Regularization techniques including dropout and L2 regularization are implemented to prevent overfitting. Loss appeared to converge during training and the last model weights were saved for inference. In order to compare our samples to those obtained by MD, we performed model inference across the entire test set. Inference is performed by providing the first frame of the MD simulation and running the reverse diffusion process for a given number of steps (here 20) to obtain predicted output frames. After post-processing, this resulted in 100 protein-ligand frames for each system, allowing for direct comparison to the 100 frames obtained via MD.

### 3.5 Metrics and Evaluation

To evaluate our model, we performed inference for the entire test set and computed qualitative and quantitative metrics. The first metric was to compare the configuration spaces sampled from MD to those sampled by our model. To this end we performed principal component analysis (PCA) directly on the coordinates of the poses obtained from MD. Using these computed PCA loadings, we transformed the samples generated by our model and plotted the first two principles components. The second metric we employ is a quantitative measure on the coverage our samples provide. This comprehensive evaluation metric is based on the pairwise Root Mean Square Deviation (RMSD) between generated poses and the set of MD molecules:

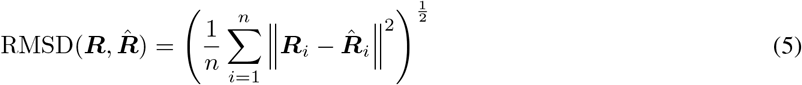

Specifically, we calculate the RMSD for each pair and then assess coverage at various threshold levels (0.5Å, 1.0Å, 1.5Å). Coverage is determined by examining each frame of the ground truth set to ensure there is at least one generated frame within the specified RMSD cutoff, capturing the model’s ability to replicate existing molecular structures accurately. Conversely, we also verify for each frame in the generated ensemble whether it corresponds to at least one frame in the ground truth set within the same cutoff values, thereby evaluating the model’s capability to produce novel, yet plausible, molecular configurations. This bidirectional assessment allows us to gauge both the completeness and the reliability of the generative model in producing molecular structures that are both diverse and representative of the true distribution.

## 4 Results and Discussion

Figure 2A shows visualized results of sampled protein-ligand states compared to those obtained via MD. Overall the visual inspection shows good agreement between the explored space, with the ligand limited to its bound pocket with some flexibility among particular atoms and groups. Figure 2A also shows the results of the PCA map that displays the configurational space of the MD and model generated frames. From these results it can be seen that the model is able to produce frames with good phase space overlap to those seen in MD. Although most ligands do not change considerably during the 10ns simulation, the observed movements seem to be consistent between the MD and generated frames. For example, in the example with PDB 1RWX, there is a flexible portion of the ligand in the middle that is sampled in two predominant conformations during the MD, while the rest of the ligand generally remains bound in the same way. The 1RWX samples obtained using our model show a similar pattern, with this flexible region of the ligand sampled in both states. Since we only have 100 frames available from the MISATO dataset, it is difficult to visualize the density of these samples. We believe the generated samples have a similar density to those from MD and are amenable to re-weighting under an energy function.

**Figure 2:**
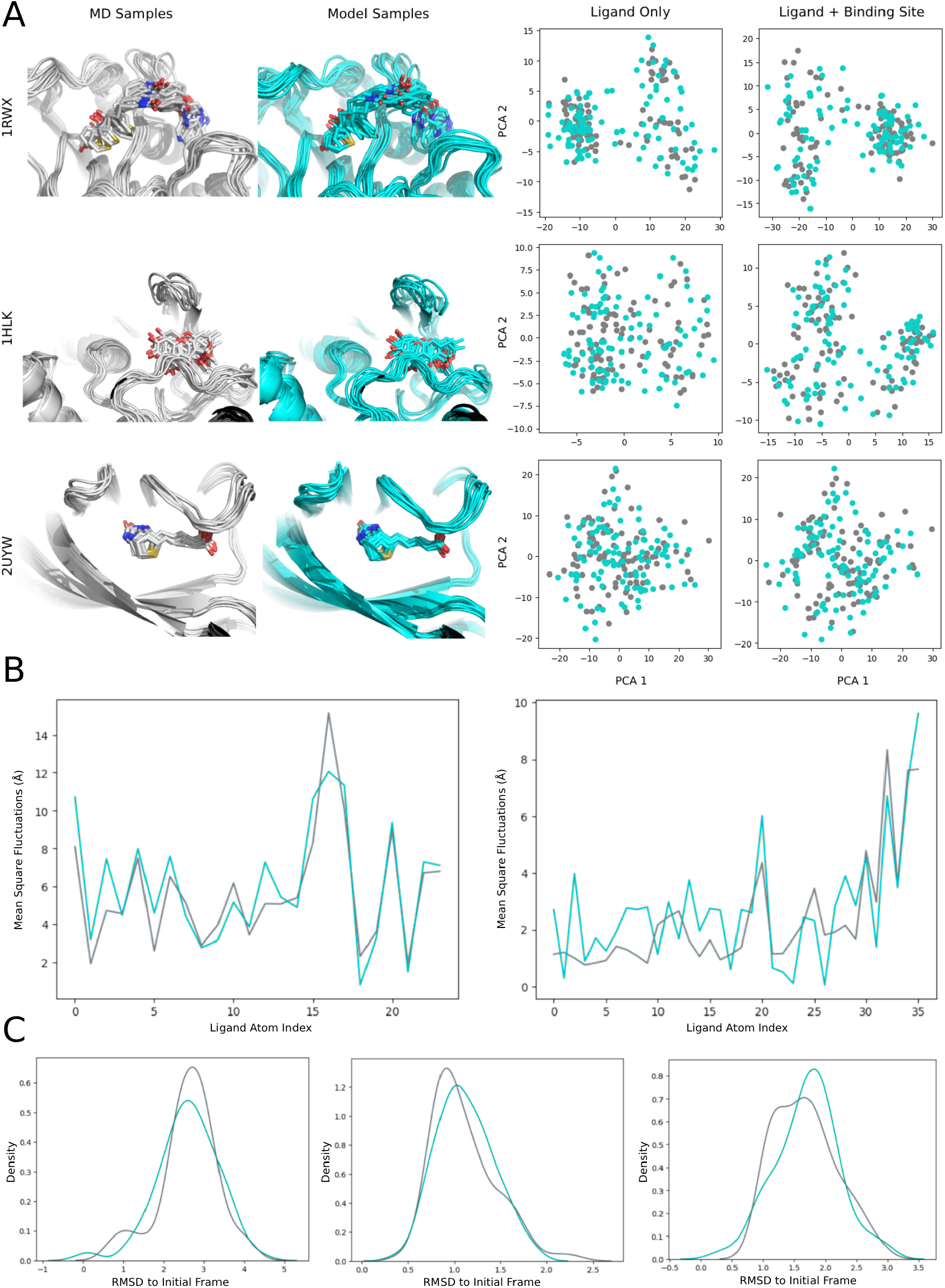
A) Visualized samples from the model (cyan) in comparison to molecular dynamics (gray). Scatter plots of the first two principle components of the ligand (and binding site) coordinates. B) Ligand mean square fluctuation plots. C) RMSD distribution KDE plots.

Figure 2B shows two examples of ligand mean square fluctuation plots. This plot demonstrates the atom specific tendency to move during the simulation and that this behavior can be replicated remarkably well by our models’ samples. Furthermore, Figure 2C shows the distribution of RMSD values with respect to the initial frame given. Once again, we see a strong agreement between the samples coming from MD and those from the model. Collectively, these results qualitatively show the capabilities of our model in accurately sampling short protein-ligand dynamics in parallel without the need for sequential molecular dynamics simulations.

Table 1 shows the results of the RMSD coverage metric described in Section 3.5. At a cutoff of 1.0 Å RMSD, we see that the model is able to recapitulate roughly 90% of the conformations from MD. Extending this to 2.0 Å RMSD covers 98% of the MD conformations. These results quantitatively confirm that our model is capable of producing samples with high overlap and coverage of those obtained via MD. This aspect is critical, especially for subsequent calculations of stability or free energy where extensive sampling and coverage of the free energy surface is necessary to produce accurate results.

**Table 1:** RMSD coverage results.

| RMSD cutoff (Å) | MD Coverage | Model Coverage |
| --- | --- | --- |
| 0.5 | 76% | 79% |
| 1.0 | 88% | 91% |
| 1.5 | 95% | 97% |
| 2.0 | 98% | 100% |

To conclude the discussion, we will mention the current limitations of this approach and routes for further research. While the model demonstrates reasonable results for the shorter length simulations it was trained and tested on, it has not yet been proven on longer length simulations that are required to observe larger conformational changes. As simulation datasets continue to grow, the possibility of adapting this model to longer length simulations is an exciting future possibility. Another limitation, as is often the case in deep learning based models, is that there is no guarantee that the samples are physically reasonable and do not contain artifacts such as steric clashes. In the current work, this is dealt with through a post-processing filter which is not ideal since it expends additional sampling time and does not represent the true Boltzmann distribution. In future works, constraints could be integrated into the diffusion process, ensuring that outputs never contain these issues and sampling can be straightforward.

## 5 Conclusion

This study successfully implemented a flow-based diffusion model to predict protein-ligand interactions from molecular dynamics (MD) simulation frames, demonstrating a strong alignment between the model-generated frames and those from MD simulations. By integrating dynamic simulation data, the model effectively captures essential conformational transitions, offering significant insights for computational drug discovery. The results affirm the model’s potential as a robust tool for understanding and predicting the dynamic processes essential for biomolecular interactions, thus advancing the predictive capabilities necessary for innovative therapeutic developments.

## Footnotes

1 3This paper is a work-in-progress and will be updated in the near future.

